# Antibacterial potential of selected extracts and silver nanoparticles from bacterial endophyte harboured by *Cola acuminata* and *Cola nitida* (Sterculiaceae) roots

**DOI:** 10.1101/2025.08.24.672022

**Authors:** Jean Baptiste Hzounda Fokou, Judith Caroline Ngo Nyobe, Germaine Dikehed Lamana, David StanyTombe Koukolin, Simone Véronique Fannang, Rosane Ngome, Philippe Belle Ebanda Kedi, Elumba Prosper Asue, Jean Emmanuel Mbosso Teinkela, François Eya’ane Meva

## Abstract

**Introduction:** Antimicrobial resistance (AMR) is currently a global health issue in most countries. Harnessing endophytic at present. Antibiotic resistance occurs when bacteria can adapt and grow in the presence of antibiotics. Endophytic microorganisms as ‘bio-factory’ of natural bioactive compounds, and their potential in nanotechnology remains largely under explored. The aim of this study was to evaluate the antibacterial activity of extracts and silver nanoparticles synthesised using bacterial endophytes isolated from *Cola acuminata* and *Cola nitida* roots.

**Methodology:** The roots of each plant were harvested, washed, cut and surface sterilised. The sterile pieces were placed on the surface of nutrient agar and incubated, after which the microorganisms were sub-cultured until pure colonies were obtained. Secondary metabolite production was then undertaken over a 12-day fermentation period in Mueller Hinton Broth, conducted under stringent aseptic conditions. Each microorganism was fermented in two different tanks. The first was used for the preparation of crude extracts, and the second for the synthesis of nanoparticles. The *in vitro* antibacterial activity was determined using the broth microdilution method against *Pseudomonas aeruginosa, Proteus mirabilis, Acinetobacter* sp. and *Escherichia coli*. The MIC and the time-kill kinetic were used to determine the inhibiting parameters. The endophytes that produced the most promising effects were identified using microscopy and MALDI-TOF techniques.

**Results:** 22 extracts were obtained from *Cola* nitida and Cola *acuminata* endophytes extracts (16 and 6 respectively. Crude extracts and silver nanoparticle). The most active material were the crude extracts from *Cola acuminata* endophytes were found to be the most active preparations. The MALDI-TOF identification method yielded the designation of NPMRU 6508, NPMRU 6511, NPMRU 7045 and NPMRU 7063 as *Bacillus cereus*. Furthermore, NPMRU6113 and NPMRU7047 were identified as *Brevibacterium sp*.

**Conclusion:** This study supports the use of endophytes derived from *Cola acuminata* and *Cola nitida* roots to combat four bacterial strains involved in the development of antibiotic resistance.

## 1. Introduction

The management of patients suffering from bacterial infectious diseases has become a major concern for medical staff due to the large number of cases of antibiotic resistance (1,2). According to the WHO, resistance occurs when bacteria can adapt and grow in the presence of antibiotics. This resistance is caused by the irrational use of antibiotics, poor patient compliance with treatment, and the variety of antibiotics used (2,3). In response, bacteria have developed sophisticated defence mechanisms, including the production of enzymes capable of inactivating antibiotics, modification of antibiotic targets, alterations in plasma membrane permeability, and the activity of efflux pumps, among other mechanisms. This escalating phenomenon has been identified as a significant public health concern by the WHO(2–4).

In view of the above, the urgent need to identify and propose new therapeutic approaches to treat these infections is becoming increasingly apparent(5).The exploitation of natural resources appears to be one of the solutions to this problem, due to their diversity and greater reserves of active substances(1,5).Plants have been a valuable source of new antibiotic compounds; however, their over-exploitation have raised concerns in terms of biomass used which could lead to flora destruction . In lieu of this, endophytes (microorganisms residing within plants) offers a viable alternative for researchers worldwide. These microorganisms, whilst residing within the plant, do not cause any harm to the plant itself (6–11), but triggered the production of molecules with a variety of additional chemical properties which utilise novel mechanisms of action against pathogenic microbes.

Nanotechnology has emerged as a particularly promising research domain due to the size-dependent properties of nanoparticles, which confer them immense versatility for medical and technological applications. This field of study focuses on materials at the nanometric scale, defined as objects measuring between 1 and 100 manometer (nm) in at least one dimension. At this scale, particles can easily cross the bacterial membrane to induce their effects (12–14). The synthesis of nanoparticles, particularly silver nanoparticles is gaining growing interest. occurs through two distinct approaches: the “top-down” method, which relies on physical principles, and the “bottom-up” approach, which utilises chemical and biological processes(12). However, physical and chemical processes are costly and generate toxic substances that are detrimental to the environment and humans. Consequently, the biological method of synthesis, known as “green synthesis”, is an inexpensive, non-toxic, and environmentally friendly process that uses micro-organisms and plant extracts as agents (13–16).

To the best of our knowledge, no studies have hitherto been carried out on silver nanoparticles obtained from *Cola acuminata* and *Cola nitida* endophytes. The present study aims to address this knowledge gap by isolating bacterial endophytes from both Cola roots, synthesising silver nanoparticles and extracts, and evaluating their antibacterial activity against some resistant strains.

## 2. Materials and Methods

### 2.1. Study characteristic and authorisations

This experimental study was carried out at the Laboratory of Pharmacology, FMSP/UDo and the Centre Pasteur du Cameroun between December 2022 and June 2023. It was conducted under research authorization number 0002/UD/FMSP/DSP and 347/UD/FMSP/DSP, granted by the Department of Pharmaceutical Sciences, Faculty of Medicine and Pharmaceutical Sciences - the University of Douala (FMSP-Udo), and was approved by ethical clearance approval number 3827 CEI-UDO/06/2023/T.

### 2.2. Biological material: plants and micro-organisms

The *Cola acumunata* and *Cola nitida* roots were harvested in the littoral region of Cameroon at Njombe-Penja and identified at the National Herbarium of Cameroon by comparison with voucher speciment under identification number 5368/SRFK and 48651 SRF Cam respectively.

The reagents and solvents of analytical grade included Mueller Hinton Broth (TM MEDIA), Agar bacteriologic and Nutrient Broth (Lyophychem), Dichloromethane (DCM), and Methanol (Solevo Cameroon Sarl), and Resazurine (SIGMA Aldrich).

The isolates were obtained from the Laboratory of Microbiology at the General Hospital of Douala, and they include *Escherichia coli, Proteus mirabilis, Acinetobacter* sp., and *Pseudomonas aeruginosa*.

### 2.3. METHODS

### Endophyte Isolation and Purification: Surface Sterilization Protocol

The initial phase of endophyte isolation involved a surface sterilization protocol for plant roots, adhering closely to the methodology previously outlined in Hzounda et al. [17].

Firstly, roots from each plant were subjected to a thorough cleansing under running tap water to remove gross debris. Subsequently, the roots were sectioned into small fragments, approximately 1 cm^2^ in size, hereafter referred to as ‘explants’. These explants were then transferred into 50 ml sterile falcons for the sterilization sequence.

The sterilization process began with a 3-minute immersion of the explants in 70% ethanol. Following this, to ensure the complete removal of any residual ethanol which could harm endophytes or interfere with subsequent steps, the explants were meticulously rinsed thrice with physiological water. A critical disinfection step involved soaking the explants in 1% sodium hypochlorite (prepared in physiological water) for one minute, immediately followed by another thorough rinse with physiological water. The final stage of surface sterilization involved drying the explants using sterile industrial paper, strictly under aseptic conditions, to prevent recontamination. The sterilization process was deemed successful and valid only if no bacterial or fungal growth was observed on the Nutrient Agar inoculated with the last rinsing water after incubation, thereby confirming the elimination of surface contaminants.

### Isolation of Endophytes from Sterilized Explants

Following successful surface sterilization, the isolation of endophytes from the prepared root explants was conducted as per the methodology established by Hzounda et al. [17].

Briefly, the sterile, dried root explants were carefully seeded onto the surface of Nutrient Agar. To specifically target bacterial endophytes and inhibit potential fungal contaminants during the initial growth phase, the Nutrient Agar was supplemented with Fluconazole at a concentration of 0.1 mg/ml. The inoculated plates were then incubated at 30°C for a period of eight days.

During this incubation period, emerging bacterial colonies were closely monitored. As colonies became visible, they were carefully harvested and repeatedly reseeded onto fresh agar plates. This iterative subculturing process was continued until uniform growth pattern, indicating pure cultutre.

Upon successful isolation and purification, the endophytes were assigned specific identification codes for traceability and future reference: isolates obtained from *Cola acuminata* were designated with the code NPMRU 70XX, while those derived from *Cola nitida* were coded as NPMRU 6XXX.

#### 2.3.1. Fermentation process

The fermentation process was caried out according to Hzounda et al. (17) with minor modification. The bacterial endophyte strains obtained after purification were inoculated (five 10ul loops) into 125 ml flasks containing 50 ml of sterile Mueller Hinton Broth. Two flasks were prepared per endophytes. These flasks were sealed, and the fermentation process was carried out for 12 days at room temperature. The flasks were hand-shaken twice a day throughout the fermentation process. Upon the completion of the 12-day fermentation period, the resulting broth from each pair of flasks was designated for specific downstream applications:

- Flask 1 (Nanoparticle Synthesis): The contents of one flask per endophyte strain were utilized directly for the biosynthesis of nanoparticles. This typically involves processes where the microbial metabolites act as reducing and capping agents for metal ions.
- Flask 2 (Crude Extract Preparations): The contents of the second flask were reserved for the preparation of crude extracts.

#### 2.3.2. Synthesis of silver nanoparticles (AgNPs) from endophytes

#### Pre-Synthetic Preparation

Following an initial incubation period during which the bacterial endophytes grow and produce metabolites, the entire bacterial endophytes mixture is subjected to a unique separation process. To facilitate the effective settling of bacterial cells, the mixture is placed in a refrigerator at a chilling temperature of –13°C. This low temperature promotes the aggregation and sedimentation of the bacterial biomass. After this cold treatment, two distinct phases are observed: a solid pellet containing the settled bacterial cells, and a clear supernatant liquid. This supernatant is rich in the secondary metabolites excreted by the endophytes, which serve as the primary reducing and capping agents for the silver ions, facilitating their transformation into stable silver nanoparticles. The pellet, consisting mainly of the bacterial cells, is discarded.

#### The Nanoparticle Synthesis Reaction

With the metabolite-rich supernatant prepared, the next phase involves the actual synthesis of the silver nanoparticles. This step is adapted from established methodologies (18) with specific modifications to optimize the biogenic process.

The purified supernatant, containing the secondary metabolic compounds, is carefully mixed with a solution of silver nitrate (AgNO3) at a concentration of 4mM. The mixed solution is then immediately transferred to a dark environment and incubated at room temperature for a period of 24 hours.

#### Confirmation of Silver Nanoparticle Formation

After the 24-hour incubation period, the solution was passed through a UV-Visible spectrophotometer, scanning a broad wavelength range, typically between 200 and 800 nm. The unequivocal sign of successful silver nanoparticle formation is the appearance of a distinct surface plasmon resonance (SPR) peak. Solutions whose spectra show this characteristic plasmon resonance within the specific wavelength range of 390 and 550 nm were confirmed to contain silver nanoparticles.

#### 4. Purification and Concentration Determination of AgNPs

To separate the formed nanoparticles from any residual impurities, unreacted silver ions, and components of the culture medium, the solutions showing clear plasmon resonance are subjected to centrifugation. This process is carried out at 6500 rpm for 20 minutes. supernatant containing impurities were carefully decanted. To further enhance the purity of the nanoparticles, the obtained pellet is subjected to a rigorous washing procedure: it is washed 3 times with distilled water. After washing, the purified nanoparticles are re-suspended in 10 mL of distilled water. To determine the precise concentration of the synthesized nanoparticles, 1 mL aliquots of the thoroughly mixed AgNP suspension are collected and placed in an oven and completely dried. The dried nanoparticle mass is then accurately weighed. This provides the mass of nanoparticles per milliliter, allowing for a precise calculation of the nanoparticle concentration in the stock suspension.

#### 2.3.3. Optimisation of nanoparticle synthesis parameters(14)

The reaction parameters such as AgNO3 concentration, pH, temperature, reaction time, temperature, reaction time and volume ratio of extract and AgNO3 were optimised to achieve maximum nanoparticle synthesis. 1 mM, 2 mM, and 4 mM silver nitrate solutions were prepared by diluting a 50 mM stock solution with distilled water. The prepared solutions were then stored in a cool, dark place in the laboratory.

Effect of pH: pH plays a crucial role in the synthesis process. Optimization was performed at pH values of 6.3, 8, 10, and 12, adjusted using a KOH solution at room temperature (25°C). The synthesis was carried out with 2 mM AgNO3 at a 1:1 volume ratio and an incubation time of 24 hours.

Effect of Silver Nitrate Concentration: The impact of different AgNO3 concentrations (1 mM, 2 mM, and 4 mM) on the optimized parameters was evaluated under the conditions of pH 12, a 1:1 volume ratio, room temperature (25°C), and a 24-hour incubation period. A noticeable transition was observed between 2 mM and 4 mM AgNO3.

Effect of Temperature: To assess the influence of temperature, the reaction was conducted at 25°C, 40°C, 60°C, and 80°C while maintaining a 1:1 volume ratio, pH 12, 4 mM AgNO3 concentration, and a 24-hour incubation period.

##### Nanoparticle characterization

Nanoparticle synthesis was characterized by Ultraviolet visible spectroscopy as describe earlier(19). Briefly, the formation of nanoparticles begins immediately after contact between plant extract and AgNO3 solution. The colour of the mixture of plant extract and AgNO3 changes from clear to yellow and dark brown due to the formation of plasmons at the colloid surface, thus indicating the synthesis and growth of silver nanoparticles. This later Plasmon can by monitored by UV-vis scan from 200 to 900 nm on a spectrophotometer.

##### Extract preparation

The active metabolites of the endophytes were extracted using two organic solvents, methanol (MeOH) and dichloromethane (DCM)(17). Following a 12-day incubation period, the fermentation process was terminated by the addition of 50 ml of methanol. The flasks, which now contained 100 ml of solution, were homogenised and left to stand for 48 hours. The solution obtained was then mixed in an Erlenmeyer flask with 100ml of DCM, homogenised, and introduced into a separating funnel to separate the two phases obtained. The organic phase containing the secondary metabolites was recovered and left at room temperature until total evaporation of the DCM and drying of the extract. The extracts obtained were designated E45, E47, E48, E49, E62 and E63, and the extraction yield of the bioactive metabolites was calculated by the weight/volume (W/V) x100 method.

#### 2.3.4. Evaluation of the antimicrobial activity of silver nanoparticles and extracts

The methodology employed in this study was micro-dilution in a medium as delineated by the Clinical and Laboratory Standards Institute (CLSI) in 2020(20).

##### Preparation of the bacterial inoculum

The bacterial isolates to be tested were activated on Mueller Hinton agar and then incubated for 24 hours at 37°C. Following this, two to three colonies were selected and introduced into tubes containing a 10ml solution of sterile NaCl. These were then compared with a chemical solution of 0.5 McFarland turbidity (consisting of 0.05ml 1% BaCl2 and 9.95ml 1% H2SO4%). The solution was then adjusted to a turbidity level comparable to that of the chemical solution, with the objective of obtaining a bacterial suspension of 0.5 McFarland or 1.5 x 10^8^ CFU/mL. Subsequently, the bacterial solutions were diluted with Mueller-Hinton broth to achieve a concentration of 3 x 10^6^ CFU/mLprior to utilisation.

##### Single concentration testing of nanoparticle and extracts

The test was conducted as follows: solutions of silver nanoparticles (AgNPs) and extract were prepared at 5 mg/ml and 10 mg/ml, respectively, by dilution in 20% DMSO. The test was carried out on the nanoparticles and the extract at concentrations of 100µg/ml and 200µg/ml, respectively, on isolates *Escherichia coli, Proteus mirabilis, Actinobacter sp*., and *Pseudomonas aeruginosa*. The cell viability was indicated by 10 µl of resazurin(21). Ceftriazone was used as a positive control at 200 µg/ml.20 µl of the solutions were transferred from the mother plate to the test plate. Then 30 µl of Mueller Hinton Broth culture medium was added, 50 µl of inoculum with a load of 3x10^6^ CFU/ml for each microorganism was added and the plate was incubated for 24 hours at 37°C. 20h after incubation, 10 µL of resazurin(0,5mg/ml) was added to each well, and the plates were incubated for 4h. The change in colour from blue to pink indicated bacterial growth, and this was noted by the sign (0) sign. The change in colour from blue to violet indicated partial inhibition of pathogenic isolates, and this was noted by the sign (M). The persistence of the blue colour of the resazurin indicated total inhibition of pathogenic strains, and this was noted by the (1) sign.

##### Determination of the minimum inhibitory concentration (MIC)

The minimum inhibitory concentration (MIC) is the lowest concentration of the antimicrobial agent that inhibits all visible growth of a bacterial strain after 18-24 hours of culture at 37°C(20,21). The test was performed using the same protocol describe for the single concentration testing with minor adjustments. The adjustment, consist of the serial dilution of the extracts and nanoparticle that highlight the total inhibition. The concentrations series were 100, 50, 25, 12.5 and 6.5, 3.125µg/ml and extract 200, 100, 50, 25, 12.5, 6.5 and 3.125µg/ml for nanoparticle and extract respectively. The tests were done in triplicate but the resazurin was added to wells and the third was used for the MBC determination. The MIC corresponds to the well with the lowest concentration that retained the blue colour of the resazurin.

##### Determination of minimum bactericidal concentration (MBC)

The minimum bactericidal concentration (MBC)was performed by inoculating 50 µL of all the wells with total inhibition into 150 µL of MHB. The plate was incubated at 37°C for 24 hours. Resazurin detection was performed as described above. The lowest concentration at which there was no change in resazurin was considered to be the CMB.

##### Evaluation of the CMB/CMI ratio

The calculation of the CMB/CMI ratio is used to confirm the bacteriostatic or bactericidal nature of a substance. If the ratio is greater than or equal to 4, the substance is considered to be bacteriostatic; if it is less than 4, it is considered to be bactericidal; if it is equal to 1, it is considered to be completely bactericidal.

#### 2.3.5. Time Kill Kinetic

This test was performed as previously describe(22). With minor adjustment. Briefly, MIC/2, 1× MIC and 2× MIC of Nanoparticle and extracts were considered, bacterial inoculum of 3×106 CFU/ml of *Proteus mirabilis, Acinetobacter sp* and *Pseudomonas aeruginosa* isolates. The test procedure was similar to that of the MIC determination with the adjustment on the final volume that was 10ml here. At T0, T2h, T4h, T6h, T8h, T12h to T24h incubation, an 1ml aliquot was collected and 100 µL was added and incubated for up to four hours. The log [CFU/ml] versus time graph was plotted.

#### 2.3.6. Characterisation of the endophytic bacteria with the highest antibacterial activity

The endophytes with targeted potential were subjected to macroscopic identification by visual observation of the shape, size, colour and appearance of the colonies, microscopic identification by optical microscopy of the shape, appearance and mobility of the bacteria, and identification by matrix-assisted laser desorption/ionisation-time of flight (MALDI-TOF) mass spectrometry.

##### MALDI TOF identification

The MALDI-TOF based identification of the bacteria were done as previously describe (23) briefly, 24h old colonies were transferred using a seeding loop to a dot of a 96-well stainless-steel target plate. Plate extraction was performed by covering each spot with 1 µl of 70% formic acid solution and air drying for 2 min. 1 µl of CHCA (a-cyano-4-hydroxycinnamic acid) matrix in 50% acetonitrile and 2.5 trifluoroacetic acid was then applied to each sample and air dried. Measurements were made on a mass spectrometer using software. The positives were extracted with an acceleration voltage of 20 kV in linear mode. The spectra were analysed for statistically significant masses. The spectrum was imported into the Biotyper software and the generated biomarker spectrum for each bacterium was compared with reference spectra.

##### Statistical analysis

Microsoft Office Excel Professional Plus 2019 was used for data entry and calculation of means and proportions.

GraphPad Prism 8.0.1 was used for variance analysis and curve plotting, employing a one-way ANOVA test with a significance threshold of P < 0.05.

## 3. Results and discussion

### 3.1. Results

#### 3.1.1. The isolation and purification of endophytes

Following the isolation and purification processes, a total of 22 bacterial endophytes were obtained, distributed as follows: six bacterial endophytes were obtained from *Cola Acuminata* roots NPMRU7045, NPMRU7047, NPMRU7048, NPMRU7049, NPMRU7062 and NPMRU7063, while five were obtained from *Cola nitida* roots bark and name NPMRU 6501, NPMRU 6130, NPMRU 6133, NPMRU 6134, NPMRU 6135 and eleven from *Cola nitida* root NPMRU6504, NPMRU6502., NPMRU6503, NPMRU6505, NPMRU6136, NPMRU6506, NPMRU6508, NPMRU6509, NPMRU6510, NPMRU6511, NPMRU6512.

#### 3.1.2. Synthesis of Nanoparticles

The synthesis of AgNPs from endophytes led to a colour change upon the addition of AgNO3 (Figure 1A and B), while the control group showed no colour change. UV-Visible spectroscopic measurements of the solution were obtained in the range of 350-550nm, which represents surface plasmon resonance, and a maximum absorbance peak was obtained at approximately 412nm (figure 1 C).

**Figure 1:**
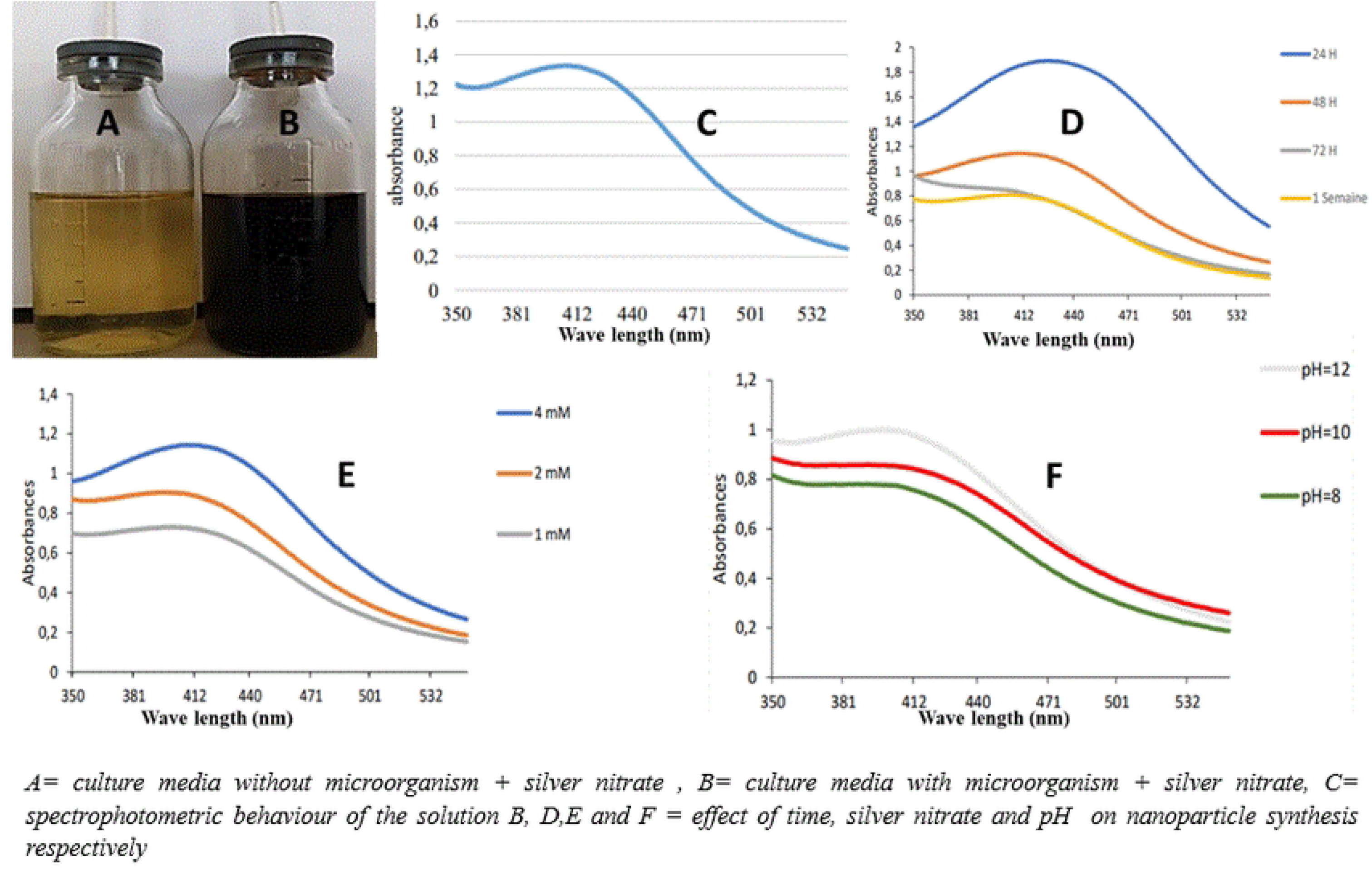
visual and spectrophotometric behaviour of the nanoparticle synthesis

The results of AgNPs concentration, the incubation time and the pH are presented on figure 1 D, E and F. From this figure, stable maximum nanoparticles are obtained at 24 hours relatively to the volume of solution used. At increased incubation time the plasmon resonance declines due to precipitation at the bottom of the flash (Figure 1D). The synthesis of nanoparticle increases along with the concentration NPsAg up to 4mM. The nanoparticle synthesis increases with the pH (figure 1 F).

### 3.2. Antibacterial activity

#### 3.2.1. Pharmacological screening

The endophytes produced a total of 22 nanoparticles and 22 extracts, which were then screened at concentrations of 200 µg/mL and 400 µg/mL, respectively. The results obtained are presented in the table 1 below. From that table most of the extracts and nanoparticles were inactive on the tested isolate at the tested concentrations. However, the extracts from endophyte NPMRU 7047 showed the best effect as it inhibited all the bacteria strains. As well the nanoparticles from the same endophytes showed a moderate activity on all the tested strains. Extracts and nanoparticle from NPMRU6133 and nanoparticles from NPMRU 6511 were active on *Pseudomonas aeruginosa* and Acinetobacter sp. Nanoparticles from NPMRU6508 were active on *Pseudomonas aeruginosa* and *Proteus mirabilis*. NP35, NP09, NP45, and NP63 were active selectively on *Pseudomonas aeruginosa. Lastly*, NP03, NP34, E34, NP06 were active selectively on Acinetobacter sp. All the other tested sampled were slightly or not active on the tested strains at the tested concentration.

**Table 1:** Result of the single dose screening of the endophyte extract and nanoparticle.

| ENDOPHYTE | PRODUCT CODE | *Pa* | *Pm* | *A* |  |
| --- | --- | --- | --- | --- | --- |
| NPMRU6501 | NP01 | 0 | 0 | 0 | 0 |
|  | E01 | 0 | 0 | M | 0 |
| NPMRU6130 | NP30 | 0 | 0 | M | 0 |
|  | E30 | 0 | 0 | 0 | 0 |
| NPMRU6502 | NP02 | M | 0 | M | 0 |
|  | E02 | 0 | 0 | 0 | 0 |
| NPMRU6133 | NP33 | 1 | 0 | 1 | 0 |
|  | E33 | 1 | 0 | 1 | 0 |
| NPMRU6503 | NP03 | 0 | 0 | 1 | 0 |
|  | E03 | 0 | 0 | 0 | 0 |
| NPMRU6134 | NP34 | 0 | 0 | 1 | 0 |
|  | E34 | 0 | 0 | 1 | 0 |
| NPMRU6504 | NP04 | 0 | 0 | 1 | 0 |
|  | E04 | 0 | 0 | 0 | 0 |
| NPMRU6135 | NP35 | 1 | 0 | 0 | 0 |
|  | E35 | 0 | 0 | 0 | 0 |
| NPMRU6506 | NP06 | 0 | 0 | 1 | 0 |
|  | E06 | 0 | 0 | 0 | 0 |
| NPMRU6136 | NP36 | 0 | 0 | 0 | 0 |
|  | E36 | 0 | 0 | 0 | 0 |
| NPMRU6508 | NP08 | 1 | 1 | M | 0 |
|  | E08 | 0 | 0 | M | 0 |
| NPMRU6509 | NP09 | 1 | 0 | 0 | 0 |
|  | E09 | 0 | 0 | 0 | 0 |
| NPMRU6510 | NP10 | 0 | 0 | 0 | 0 |
|  | E10 | 0 | 0 | M | 0 |
| NPMRU6511 | NP11 | 1 | 0 | 1 | 0 |
|  | E11 | 0 | 0 | 0 | 0 |
| NPMRU6512 | NP12 | 0 | 0 | 0 | 0 |
|  | E12 | 0 | 0 | 0 | 0 |
| NPMRU6505 | NP05 | 0 | 0 | 0 | 0 |
|  | E05 | 0 | 0 | 0 | 0 |
| NPMRU7045 | NP45 | 1 | 0 | 0 | 0 |
|  | E45 | 0 | 0 | 0 | 0 |
| NPMRU7047 | NP47 | M | M | M | M |
|  | E47 | 1 | 1 | 1 | 1 |
| NPMRU7048 | NP48 | 0 | 0 | 0 | 0 |
|  | E48 | 0 | M | M | M |
| NPMRU7049 | NP49 | 0 | M | M | M |
|  | E49 | 0 | M | 0 | 0 |
| NPMRU7062 | NP62 | 0 | 0 | 0 | M |
|  | E62 | 0 | M | M | 0 |
| NPMRU7063 | NP63 | 1 | M | M | 0 |
|  | E63 | 0 | 0 | 0 | 0 |

Pa= *Pseudomonas aeruginosa*, Pm= *Proteus mirabilis* A=*Acinetobacter* sp ; Ec= *Escherichia coli* ; 0= no effect, 1= growth inhibition, M= moderate inhibition, NP= nanoparticle, E= extract

From the above mentioned observations, N33, 11, 08 N45, 63, and E47 were selected for further MIC and MBC determination. E33 was not included because the sample was too small for the next stage.

#### 3.2.2. Determination of MIC and MBC of nanoparticles and extracts

The selected extracts were tested for the determinations of their inhibitory parameter (MIC and MBC). The results obtained are depicted in table 2 below. From this table, it came be observed that pseudomonas aeruginosa is the most sensitive strains towards the sample. the most active sample was the extracts from endophyte NPMRU 7047 that exhibit MIC ranging from 3.125 to 50 ug/ml. as the MBC/MIC ration were all lower than 4, the extracts was classified as bactericidal on all the tested strains. All the except Acinetobacter that were sensitive to ceftriaxone, all the other bacterial strains wee resistant.

**Table 2:**
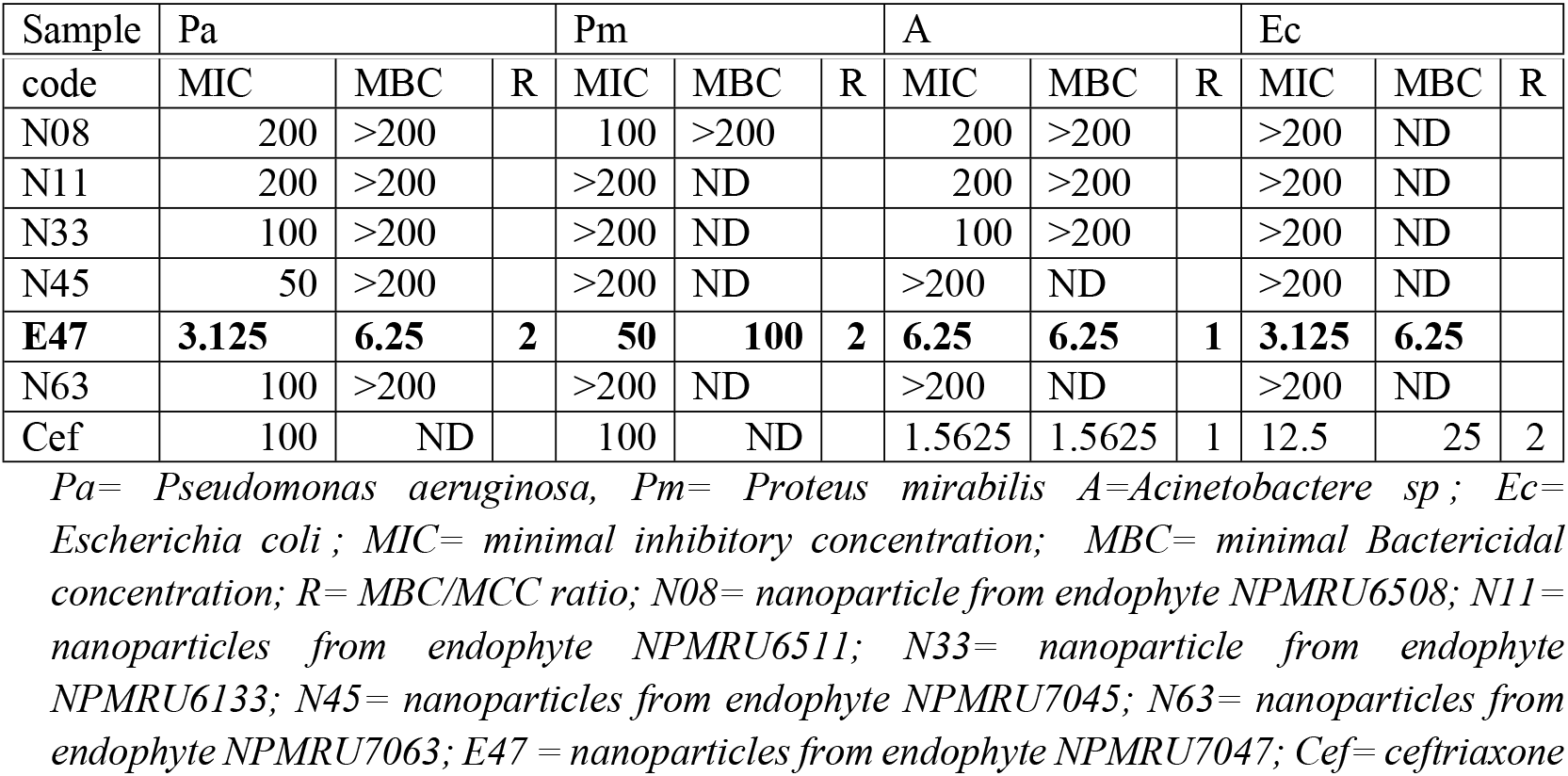
Antibacterial parameters of endophytic extracts and nanoparticle.

Based on the above-mentioned reasons, E47 and N45 were selected for time kill kinetic analysis as they were the most active extracts on *Pseudomonas aeruginosa* as the most sensitive strains.

#### 3.2.3. Time kill kinetics

Time kill kinetics were performed at different concentrations MIC/2, MIC, 2 MIC of N45, N63 and E47 against P. aeruginosa. The figures below show log [CFU/ml] vs time.

From the figure 2, statistical analysis revealed a highly significant activity of the extract in the MIC/2, MIC and 2MIC compared to the negative control (P<0.05). However, no significant difference was obtained between the different MIC/2, MIC and 2MIC (P>0.05). The higher inhibition was observed at T6h and T24 compared to the positive control.

**Figure 2:**
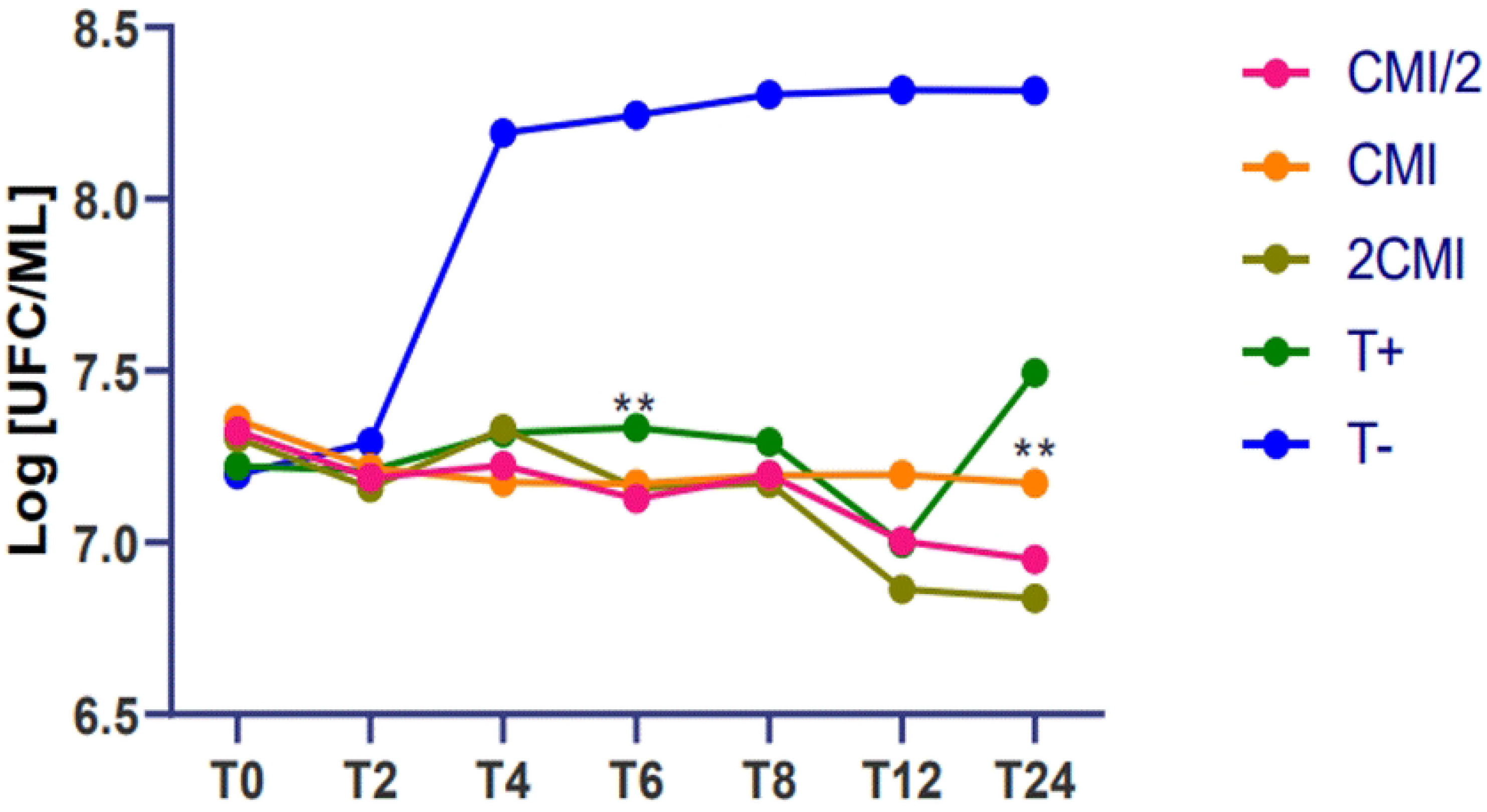
Time kill kinetic of extract from NPMRU 7047 on *Pseudomonas aeruginosa*

The time kill kinetic of the nanoparticle is presented in the figure 3 bellow

**Figure 3:**
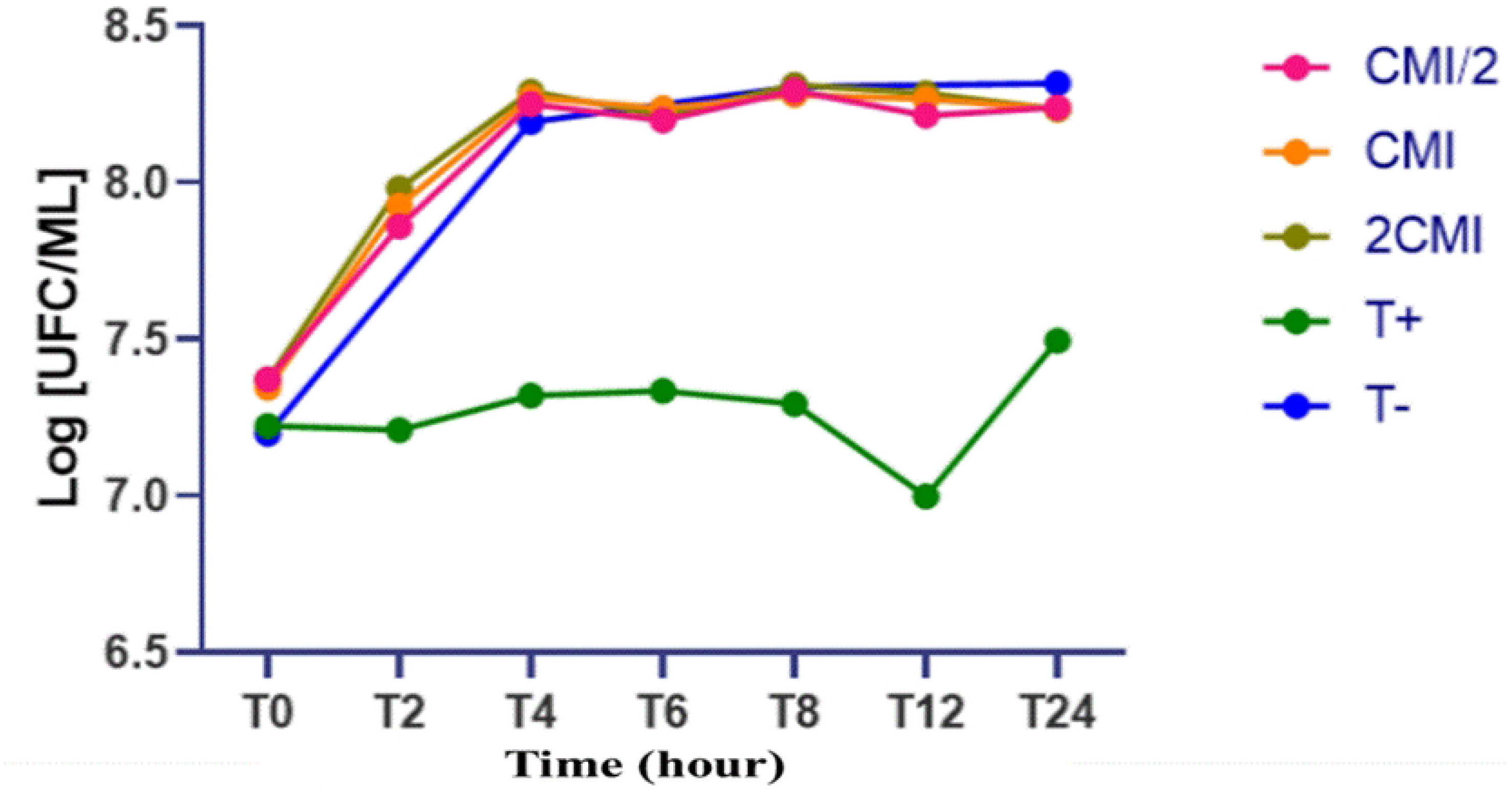
Time kill kinetic of Nanoparticle from NPMRU 7045 on *Pseudomonas aeruginosa*

From the figure 3, A non-significant activity of the N45 nanoparticle was obtained at MIC/2, MIC and 2 MIC compared to the negative control (P>0.05).

#### 3.2.4. Characterisation and identification of the most antibacterial endophytic bacteria

The endophytes NPMRU 7045, NPMRU 7063, NPMRU 7047, NPMRU6133, NPMRU 6508 and NPMRU 6511were submitted to identification due to their global effect on bacteria involved in our study.

In this vein macroscopic analysis of NPMRU 7045 showed flat, whitish, flat, whitish colonies with a creamy texture and serrated margins; NPMRU 7063 showed flat, whitish colonies with a creamy texture and semi-uniform margins and NPMRU 7047 showed pale yellow colonies with a circular shape, uniform margins and pasty texture. NPMRU 6508 and NPMRU 6511 presented flat and smooth colonies with a creamy texture, while NPMRU 6133 presented yellow colonies with a circular shape. Microscopic analysis of NPMRU7045 and NPMRU7063 were rod-shaped motile bacilli and NPMRU 7047 were motile coccobacilli. NPMRU 6508 and NPMRU 6511 appeared as rod-shaped bacilli. rod-shaped and motile. NPMRU 6133 appeared as cocci in agglomerates.

MALDI-TOF (matrix-assisted laser desorption ionisation time of flight) identification showed that the NPMRU 7045, NPMRU 7063, NPMRU6508 and NPMRU6511 identified as *Bacillus cereus* and the endophyte NPMRU 7047 and NPMRU 6133 were identified as Brevibacterium sp.

### 3.3. Discussion

This study aimed at isolating and evaluating the antibacterial effect of the extracts and nanoparticle from *Cola acuminata* and *Cola nitida* bacterial endophytes.

A total of 22 endophytes were isolated. This density of isolated endophytes may be related to the fact that the population density of endophytes is generally higher (10^5^ CFU/g) in roots than in all other plant organs(11,24). Furthermore, the population density of bacterial endophytes in roots is generally influenced by both biotic and abiotic factors. The presence of endophytes in this part of the plant confirms Strobel’s hypothesis that all plant organs have endophytes (24– 26).

The silver nanoparticles were obtained by a biosynthetic method using the supernatant (containing the metabolites) of a bacterial endophyte culture and the silver solution. The colour change is due to the reduction of Ag+ ions to AgNPs by enzymes and secondary metabolites secreted by the microorganisms(16,27). Specifically, the colour change occurs due to the vibration of free electrons within the nanoparticles surface. (12,16). The formation of nanoparticles was confirmed by obtaining UV-Vis spectroscopy of a surface plasmon resonance band between 350 and 550 nm; which is similar to the changes involving plant metabolites (28). AgNPs. The stability at 24h post reaction is due to the stabilization of the nanoparticles by the endophyte’s metabolites (12). In the reactions conditions the endophytes metabolites are able to reduce Ag+ to Ag0, and to stabilize the Ag0 obtained(29). The AgNPs formation was monitored under different conditions, varying the pH and silver nitrate concentration(12). the formation of nanoparticles increased proportionally to the different factors mentioned. Thus, basic media without microorganism favours the formation of nanoparticles and amount of available Ag+ ions increase the nanoparticles production(27). To the bes of our reading this is the first report on the synthesis of nanoparticle by bacterial endophytes isolated from *Cola nitida* and *Cola acuminata* are not known to date.

In this study, the *in vitro* antibacterial activity was evaluated against 4 bacterial isolates consisting of *Pseudomonas aeruginosa, Proteus mirabilis, Acinetobacter sp* and *Escherichia coli*. To the best of our reading, we did not find the rapport on the antibacterial effect of neither nanoparticle nor crude extracts from both *Cola nitida* and *Cola acuminata* endophytes. The characterisation of the endophytes with the greatest antibacterial activity was performed by macroscopic analysis of colonies, microscopic analysis of bacteria and MALDI-TOF analysis.t this approach is consistent with the literature where many taxa of bacteria have been successfully identified using MALDI-TOF (23,30–32). According to these analyses in one hand, four endophytes were identified as Bacillus cereus (NPMRU 7045, NPMRU 7063, NPMRU 6508 and NPMRU 6511). This is consistent with the literature where *Bacillus cereus* was identified as endophyte with plant promoting potential(33), moreover, these goes in the straight line with our finding on synthesis of nanoparticles. In fact, synthesis of nanoparticle from bacillus cereus have been documented in the literature(14,34). Furthermore, this is consistent with the antibacterial effect. Reports in the literature revealed that bacillus cereus had produced antibacterial effect (35)as well, their silver nanoparticle had antibacterial potential (14). On the other hand, two endophytes were identified as *Brevibacterium* sp. This goes in the same line with the literature that had report Brevibacterium as endophyte from many plants organs(8,36) and many other ecosystems around the word(31,37). As well, this is consistent with the silver nanoparticle synthesis(34). All this goes in the same line with the antibacterial effect of the extract and nano particle(8,18) However, the difference observed between the two Brevibacterium can be related to the plant from which they were isolated from. In fact, the same endophytes isolated from different plants or environment produce different type of secondary metabolite and hereby different pharmacological activity(8,37).

## 4. Conclusion

The present study identified a bacterial endophyte isolated from Cola acuminata and Cola nitida. The results obtained demonstrate that the silver nanoparticle synthesised from the extracellular metabolites of these endophytes exhibits poor antibacterial activity against the tested strains. However, *Brevibacterium sp*. NPMRU 7047 is a highly promising source of bioactive compounds with the potential to combat bacterial infection.

## Acknowledgement

The authors acknowledge the Dean of the Faculty medicine and pharmaceutical sciences for providing space and equipment for this project.

